# Diversity Predicts Ability of Bacterial Consortia to Mitigate a Lethal Wildlife Pathogen

**DOI:** 10.1101/123968

**Authors:** Rachael E. Antwis, Xavier A. Harrison

## Abstract

Symbiotic bacterial communities can protect their hosts from infection by pathogens. Treatment of wild individuals with protective bacteria isolated from hosts can combat the spread of emerging infectious diseases, but it is unclear whether the degree of bacterially-mediated host protection is uniform across multiple isolates of globally-distributed pathogens. Here we use the lethal amphibian fungal pathogen *Batrachochytrium dendrobatidis* as a model to investigate the traits predicting broad-scale *in vitro* inhibitory capabilities of both individual bacteria and multiple-bacterial consortia. We show that inhibition of multiple pathogen isolates is rare, with no clear phylogenetic signal at the genus level. Bacterial consortia offer stronger protection against *B. dendrobatidis* compared to single isolates, but critically this was only true for consortia containing multiple genera, and this pattern was not uniform across all *B. dendrobatidis* isolates. These novel insights have important implications for the effective design of bacterial probiotics to mitigate emerging infectious diseases.

## INTRODUCTION

The last 50 years have seen the emergence of several virulent wildlife pathogens with broad host ranges (Tompkins et al 2015). These emerging infectious disease (EIDs) have decimated wildlife populations globally, and are a major contributor to the current global loss of biodiversity (e.g. Skerratt et al 2007; McCallum 2012). Both climate change (Cohen et al 2017) and the global trade in animals (Tompkins et al 2015) are exacerbating the spread of EIDs, and broad-scale, effective treatments and/or prophylaxis for these pathogens in the wild are often lacking (Sleeman 2013; Garner et al 2016). Developing such treatments is often complicated by broad variation in genetic and phenotypic traits such as virulence exhibited by these pathogens (e.g. de Jong & Hien 2006; Schock et al 2010; Farrer et al 2011). Successful mitigation of EIDs in the wild demands that preventative or curative therapies demonstrate broad activity over as many genetic variants of the pathogen as possible, and developing mitigation strategies that satisfy this criterion remains a major outstanding research goal. Most EIDs are attributed to fungal pathogens, including *Pseudogymnoascus destructans* that causes white nose syndrome in bats, and *Batrachochytrium spp*., which causes chytridiomycosis in amphibians (Fisher et al 2012). *Batrachochytrium dendrobatidis* comprises multiple, deeply diverged lineages, and is capable of rapid evolution (Farrer et al 2011; 2013). Endemic hypovirulent lineages of *B. dendrobatidis* have been identified, including *Bd*CAPE (South Africa), *Bd*CH (Switzerland), *Bd*Brazil (Brazil) and a lineage from Japan (Goka et al 2009; Farrer et al 2011; Schloegel et al 2012; Rosenblum et al 2013; Rodriguez et al 2014), although there are cases where these have spread to other regions and are implicated in population declines in those regions (e.g. *Bd*CAPE in Mallorcan midwife toads, *Alytes muletensis;* Doddington et al 2013). The globally distributed and hypervirulent global panzootic lineage (*Bd*GPL) is the genetic lineage of *B. dendrobatidis* associated with phenomenal mass mortalities and rapid population declines of amphibians around the world, and is a major driver of the current “amphibian extinction crisis” (Fisher et al 2009; Farrer et al 2011; Olson et al 2012). Isolates within this lineage exhibit enormous and unpredictable variation in virulence, even within a single host species exposed under laboratory conditions (Farrer et al 2011; Farrer et al 2013). There is currently no cure for this disease in the wild (reviewed in Garner et al 2016), and given that amphibian communities may be host to multiple *Bd*GPL genotypes (Morgan et al 2007; Rodriguez et al 2014), and that continuous global movement of humans and wildlife transports the fungus, any prophylactic or curative treatment needs to effective against multiple *B. dendrobatidis* genotypes and isolates.

Bacterial probiotics represent a promising tool to combat major emerging fungal pathogens in the wild, including *Pseudogymnoascus destructans* (Hoyt et al 2015), *B. dendrobatidis*, and the closely related *B. salamandrivorans* (Martel et al 2013; 2014). Of these, probiotic research is currently most advanced for *B. dendrobatidis* (reviewed in Bletz et al 2013 and Rebollar et al 2016). Laboratory and field studies have shown host-associated bacterial communities (hereafter referred to as the ‘microbiome’) protect amphibians from *B. dendrobatidis* infection, and that it is possible to artificially augment the microbiome with ‘probiotic’ bacteria to improve survivorship in response to the pathogen (Bletz et al 2013; Becker et al 2015; Walke et al 2015).

To date, most *in vitro Bd* GPL challenge experiments have tested the ability of candidate probiotics to limit the growth just a single isolate of *Bd* GPL. This is problematic because the inhibitory capabilities of individual bacteria are not uniform across the variation presented by *Bd*GPL (Antwis et al 2015). Previous work has found no evidence of a phylogenetic signal in the ability of bacterial genera to inhibit a singular *Bd*GPL isolate (Becker et al 2015), but a major gap in our understanding concerns whether some bacterial genera are more likely to show broad-spectrum inhibition across a range of *Bd*GPLs, allowing a more focussed search for effective amphibian probiotics. Furthermore, both *in vivo* amphibian probiotic trials and i*n vitro* challenges focus on the application of a singular bacterial isolate to arrest the growth of *B. dendrobatidis*, yet the importance of a complex and diverse microbiome for resilience to infection has been repeatedly demonstrated across a range of host taxa (e.g. Dillon et al 2005; Matos et al 2005; Van Elsas et al 2012; Eisenhauer at el 2013). A novel alternative strategy involves a ‘bacterial consortium’ approach to probiotics, whereby multiple inhibitory bacterial isolates are applied simultaneously. Multi-species consortia can increase the inhibition of *Bd*GPL (Piova-Scott et al 2017), and so may offer greater inhibitory capabilities across a wider range of *B. dendrobatidis* isolates, however the generality of this pattern across multiple pathogen variants remains untested. Addressing the shortfall in our understanding is critical for developing effective tools for the mitigation of EIDs in the wild.

Here we extend previous work to quantify the ability of metabolites from both individual bacteria and co-cultured bacterial consortia to demonstrate broad-scale inhibition across a panel of *B. dendrobatidis* isolates. First, we test 58 bacterial isolates from 10 genera for inhibition against a suite of 10 different *Bd*GPL isolates to quantify; i) variation among bacterial genera in ability to demonstrate broad-spectrum *Bd*GPL inhibition; and ii) variation among *Bd*GPL isolates in susceptibility to inhibition. Second, we quantify the relative efficacy of using single bacterial isolates or bacterial consortia to modify *B. dendrobatidis* growth rates *in vitro*. Specifically, we investigate; iii) whether consortia yield stronger inhibition than single bacteria across three *B. dendrobatidis* isolates from two lineages (*Bd*GPL and *Bd*CAPE); and iv) whether the diversity of a bacterial consortium (number of member genera) affects inhibitory capabilities.

## METHODS

### Phylogeny screening

*In vitro* challenges were conducted for 58 bacteria isolated from wild *Agalychnis spp.* frogs in Belize (Antwis et al 2015) to screen for inhibitory capabilities against 10 *Bd*GPL isolates (Table 1, Figure 1). Bacteria belonged to 10 genera, with 3-11 bacterial isolates per genus (Table S1). Bacteria were previously identified (Antwis et al 2015) using colony PCR with primer pair 27F and 1492R for the 16S rRNA gene, which were sequenced at the University of Manchester, and then the forward and reverse sequences were aligned for each bacterium and blasted against the NCBI database (http://blast.ncbi.nlm.nih.gov/Blast.cgi). Inhibition challenges were conducted using an *in vitro* spectrophotometer assay method adapted from Bell et al. (2013), Woodhams et al (2014) and Becker et al (2015). Bacteria were grown by adding 50ul of frozen stock bacteria (stored in 30% glycerol, 70% tryptone solution at -80**°**C) to 15ml of 1% tryptone, and incubating at 18**°**C for 36 hours until turbid (three cultures per bacterial isolate). Although cell density has been shown to influence metabolite production in culture (Yasumiba et al 2015), we decided not to count and adjust cell density prior to inhibition trials as subsequent addition of media may alter the metabolite profiles already produced by cultures. In addition, cultures were not grown in the presence of *B. dendrobatidis* as multiple *B. dendrobatidis* isolates were tested in this study and this would have confounded results. Turbid cultures were filtered through a 0.22um sterile filter (Millipore, Ireland) to remove live cells, leaving only bacterial metabolites in solution. These were then combined across the three cultures for a given bacterial isolate, and kept on ice until *B. dendrobatidis* challenges were conducted. *Bd*GPL (Table 1) isolates were grown in 1% tryptone broth until maximum zoospore production was observed (∼3-4 days; ∼1 x 10^6^ zoospores ml^-1^). As with bacteria, three flasks per *B. dendrobatidis* isolate were grown and then combined prior to challenges to limit flask-effect. Zoospores were separated from sporangia by filtering through 20um sterile filters (Millipore, Ireland). To conduct the spectrophotometer assays, 50ul of bacterial metabolites and 50ul of *B. dendrobatidis* suspension were pipetted into 96-well plates. Each *B. dendrobatidis*-bacteria combination was run with three repeats. Positive controls were included using 50ul 1% tryptone instead of bacterial metabolites. Negative controls were included using 50ul sterile water and 50ul of heat-treated *B. dendrobatidis* for each isolate. Plate readings were taken every 24 hours for four days using a 492nm filter.

**Figure 1.**
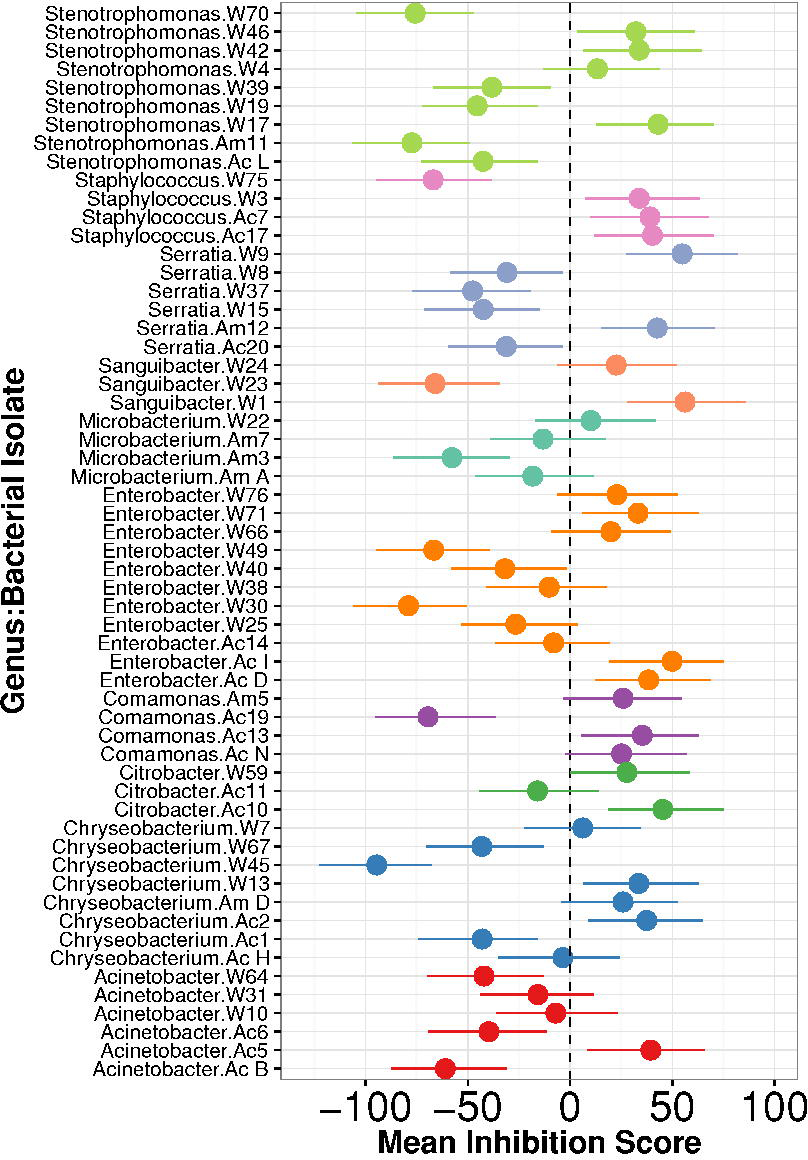
Inhibition scores of 58 bacterial strains from 10 genera when tested against 10 *Bd*GPL isolates. Estimates are derived from a Bayesian mixed effects model with bacterial isolate nested within genus, and *Bd*GPL isolate fitted as random effects. Points are conditional modes of the individual isolate random effects, marginalised with respect to *Bd*GPL isolate. Error bars are 95% credible intervals.

For each measurement, data were transformed using the equation Ln(OD/(1-OD)), and a regression analysis was used to gain the slope values for each sample over time. Slopes of triplicate replicates for each Bd/Bacteria combination were averaged, and total *B. dendrobatidis* inhibition was calculated using the formula: Inhibition (%) = [1-(slope of sample/slope of control)] x 100. A positive value represents inhibition of *B. dendrobatidis* growth, and a negative value indicates enhanced growth of *B. dendrobatidis*.

### Bacterial consortium challenges

Three bacteria were then selected from each of four genera (*Acinetobacteria, Chryseobacterium, Serratia, Stenotrophomonas*) based on their inhibition profiles; poor to medium inhibitors were selected to determine whether combining these bacteria would improve their inhibitory capabilities. Bacteria were grown individually until turbid and added to fresh tryptone either individually (strains A, B and C of each genus separately), or as a triple (strains A, B and C of each genus together to form single-genus mixes, or a random combination of strains across genera to form multi-genus consortia (Table 2)). For both individual and triple bacterial combinations, a total of 3ml of bacteria were added to 12ml of fresh 1% tryptone broth and left to grow together for 12 hours. The volume of each bacterium added depended on whether the consortium contained one or three bacteria, and the volume was split evenly between the number of bacteria added to each group. Following this, bacteria-*B. dendrobatidis* challenges were conducted using the same methods as described above against three *B. dendrobatidis* isolates (Table 1). Average inhibition percentages for each consortium-*B. dendrobatidis* combination were calculated as described above.

### Statistical Analysis

All statistical analyses were conducted in the software R v.3.3.2 (R Core Team 2016; Supplementary Information).

#### Phylogeny Data

To quantify differences among genera in proportion of *Bd*GPL isolates inhibited (inhibition score >0), we fitted a Binomial GLM with the proportion of the 10 *Bd*GPL isolates each bacterial isolate inhibited as the response, and genus as a fixed effect. We used the quasibinomial error structure as the model was overdispersed (dispersion 6.4), and tested the model containing a genus term with the reduced intercept-only model using a likelihood ratio test.

To quantify differences among genera in the *degree* of inhibition (size of inhibition score), we fitted a hierarchical model in the R package *MCMCglmm* (Hadfield 2010) with the individual inhibition scores of each bacterial isolate (n=58) for each *Bd*GPL isolate (n=10; total n = 580) as a Gaussian response. We fitted both *Bd*GPL isolate, and bacterial strain ID nested within bacterial genus as random effects. We use uninformative, parameter-expanded priors for the random effects as detailed in Hadfield (2010). We ran models for a total of 100,000 iterations following a burn-in of 10,000 iterations and using a thinning interval of 50. Posterior model checks indicated no significant autocorrelation within chains (all values < 0.05) and adequate convergence using the Geweke diagnostic (Geweke 1992). Inspection of model residuals from the frequentist analogue of this model fitted in *lme4* (Bates et al 2015) revealed normally-distributed residuals and no evidence of heteroscedasticity. Rerunning models with stronger priors has no effect on model results.

To calculate % variance in inhibition explained by *Bd*GPL isolate, bacterial genus, and bacterial strain respectively, we extracted the variance components from the variance-covariance matrix of the model above. We expressed the variance of a component *V* as a percentage of the total variance calculated as (V_BdGPL_ + V_genus_ + V_strain_ + V_residual_). We calculated both mean and 95% credible intervals using the posterior samples from the model. To construct Figs. 1 and 2, we extracted the marginal means and 95% credible intervals for each bacterial strain and *Bd*GPL isolate, respectively. That is, the bacterial strain modes are marginalised with respect to *Bd*GPL and vice versa, to quantify whether the *average* scores for each *Bd*GPL or bacterial isolate are significantly different from zero.

**Figure 2.**
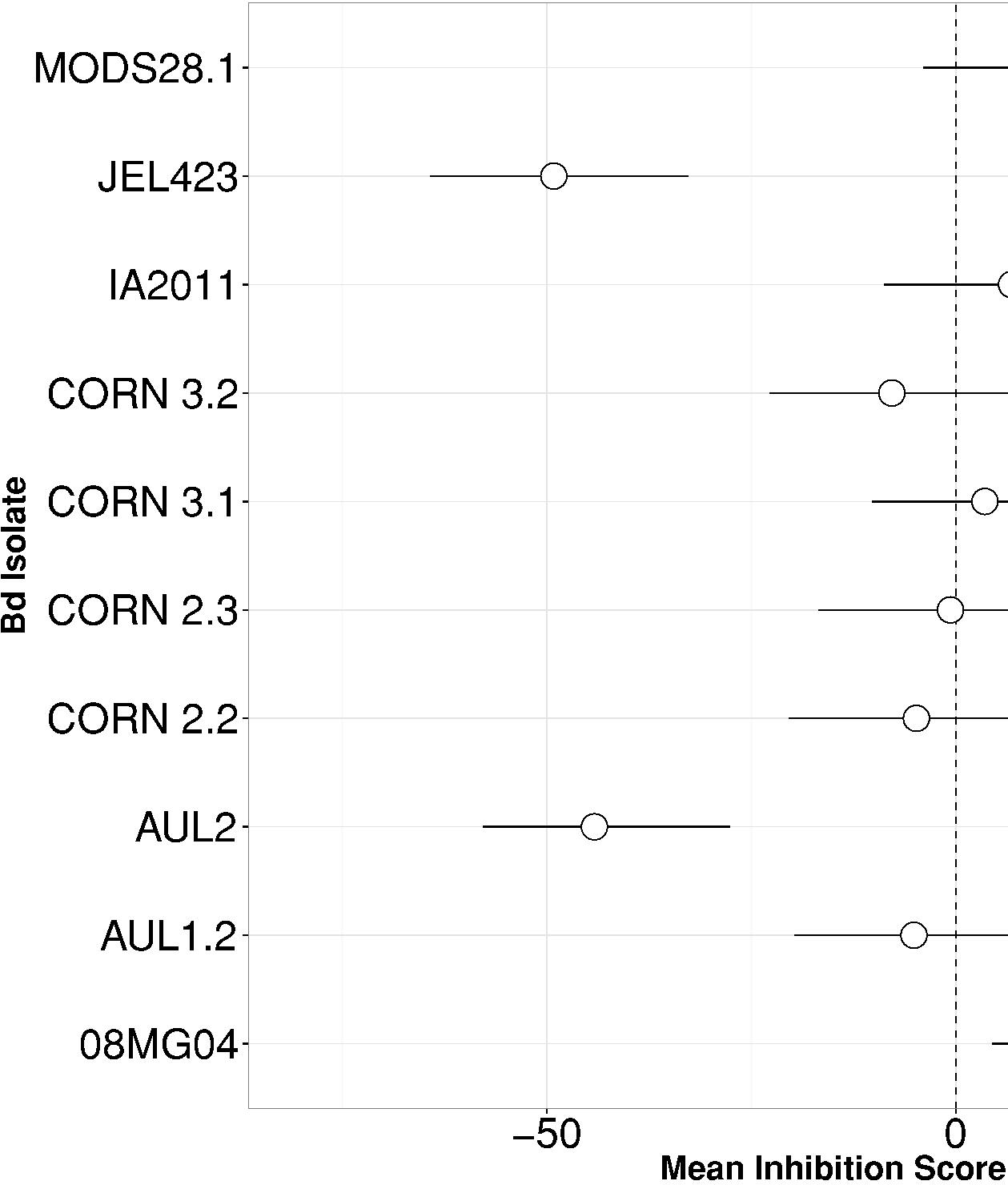
Inhibition scores of 10 *Bd*GPL isolates. Estimates are derived from a Bayesian mixed effects model with bacterial isolate nested within genus, and *Bd*GPL isolate fitted as random effects. Points are conditional modes of the individual *Bd*GPL isolate random effects, marginalised with respect to bacterial isolate. Error bars are 95% credible intervals.

#### Consortium Data

To calculate the relative mean inhibition of single-genus (SG) vs multi-genus (MG) consortia, we fitted a mixed model in *MCMCglmm* with inhibition as a Gaussian response, consortium type as a 2-level factor, and a random effect of *B. dendrobatidis* and using uninformative priors. To calculate whether consortia exhibited stronger inhibition than the mean of their individual isolates, we constructed a binary variable with outcome 1 if a consortium’s inhibition was greater than the single isolate mean and 0 if equal to or lower. We fitted this as a response in a binary GLMM with consortium type as a fixed effect, *B. dendrobatidis* as a random effect and using uninformative priors. Neither model exhibited signs of autocorrelation and Geweke statistics for both models indicated convergence.

#### In silico Probiotic Consortium Trials

To probe the relative effectiveness of single bacteria, SG consortia and MG consortia (hereafter ‘probiotic types’) for modifying the growth rates of *B. dendrobatidis*, we ran three sets of simulations, each comprising 1000 iterations. For each set of simulations, we calculated i) the proportion of times a MG consortium yielded higher inhibition than a SG consortium; ii) the proportion of times a MG consortium yielded higher inhibition than a single bacterial isolate; iii) the probability that a MG, SG or single bacterial isolate would yield at least 50% inhibition, which we class as strong inhibition. Adopting a Monte Carlo Integration approach allows us to investigate the performance of different probiotic strategies for individual *Bd* isolates. Calculating group means of each probiotic type does not allow one to calculate the frequency that one probiotic type might outperform another, as this approach does not explicitly make pairwise comparisons and in fact loses information by comparing group means. Group means can be skewed by large individual values, and therefore be misleading with respect to the efficacy of a particular strategy if the mean of that group is not reflective of the true variance in the data. However we report group means alongside these statistics where appropriate for comparison. We derived 95% confidence intervals for each test statistic by performing 10,000 bootstrap samples with replacement from the test distributions. The three simulations were as follows:

1. Averaged over all *B. dendrobatidis* isolates: For each iteration, we randomly selected a *B. dendrobatidis* isolate, and then randomly selected both a SG and a MG consortium. A Single bacterial isolate score was then selected randomly from one of the members of the MG consortium.
2. *B. dendrobatidis* specific scores: To investigate the potential for the effectiveness of consortia to differ depending on *B. dendrobatidis* isolate, we repeated the simulations as in (1) but performed 1000 simulations for *each B. dendrobatidis* isolate.
3. Sequential *B. dendrobatidis* exposure: Finally, we examined the ability of the three probiotic types to inhibit two *B. dendrobatidis* isolates encountered in series by randomly selecting two of the three *B. dendrobatidis* isolates. We assumed that the two isolates are not encountered simultaneously, as co-occurrence of two *Bd* isolates may modify their growth rates and/or a bacterial isolate’s ability to inhibit them. For each iteration, we selected a random MG and SG consortium, followed by a randomly-selected single isolate member from the MG consortium. Individual inhibition scores for these three groups were then extracted for both selected *B.* dendrobatidis isolates (i.e. probiotic ID was kept consistent over both pathogen isolates). We calculated the probability that the MG consortium would yield superior inhibition to the SG consortium and single bacterial isolate across both *B. dendrobatidis* isolates, and the probability that all three probiotic types would yield >50% inhibition.

## RESULTS

### Phylogenetic Signals of *Bd*GPL Inhibition

We assayed the ability of 58 bacterial isolates from 10 genera to modify the growth rates of 10 *Bd*GPL isolates. Mean inhibition scores ranged from100% (complete inhibition of growth) to -225% (strong facilitation of growth). At the genus level, there was no significant variation among genera in mean proportion of *Bd*GPL isolates inhibited (Binomial GLM; *χ*^2^_9_ = 6.2, p=0.72; Table 3). Six isolates from five genera showed at least weak inhibition across all 10 *Bd*GPLs, whilst seven isolates from five genera facilitated the growth of all 10 isolates (Supplementary Table S1).

Variance component analysis revealed considerably more variation in inhibition scores among bacterial strains *within* genera than among genera themselves (Fig. 1). Variation among bacterial strains within genera explained 51% [95% credible interval (CRI) 37-63%] of the variation in *Bd*GPL inhibition scores compared to just 1.3% [0.09-6%] for bacterial genus. *Bd*GPL isolate explained 15.6% [4.8-30%] of the variation in inhibition scores and highlighted two isolates whose marginal effect sizes were significantly negative (JEL423 and AUL2), and one isolate with a significantly positive marginalised inhibition score (08MG04; Fig. 2). JEL423 and AUL2 therefore exhibit strongly enhanced growth in the presence of bacterial metabolites, whereas 08MG04 is particularly susceptible to inhibition of growth. The remaining seven *Bd*GPL isolates demonstrated no evidence of systematic susceptibility to inhibition of their growth rates across the bacteria tested (Fig. 2).

### Multi-Isolate Consortia as Tools for Pathogen Mitigation

Consortia containing isolates from Multi-Genus (MG) exhibited significantly higher mean inhibition scores compared to Single-Genus (SG) consortia when marginalising with respect to *B. dendrobatidis* isolate (MG consortia mean inhibition: 36.88%; SG consortia mean: 16.9%; 95% CRI of difference 4.12 – 36.52%, p_MCMC_ = 0.02; Fig. 3). If the ability of a consortium to inhibit *B. dendrobatidis* was simply an additive function of the inhibitory capabilities of the individual bacteria it comprised, we would expect the consortium’s inhibition score to be equal to the mean of the individual inhibition scores, weighted by relative abundance. Inhibition scores of consortia greater than the mean of individual isolate scores are indicative of synergistic effects, whereby the combined pool of metabolites from multiple bacteria inhibits *B. dendrobatidis* more strongly than the individual isolates. MG consortia had a 61% probability of demonstrating stronger inhibition than the mean of their single composite bacterial isolates, which was significantly higher than the corresponding probability for SG isolates (26.6%, Mean difference 39.4% [95% Credible Interval 11.2-65.1%], p_MCMC_ = 0.01).

**Figure 3.**
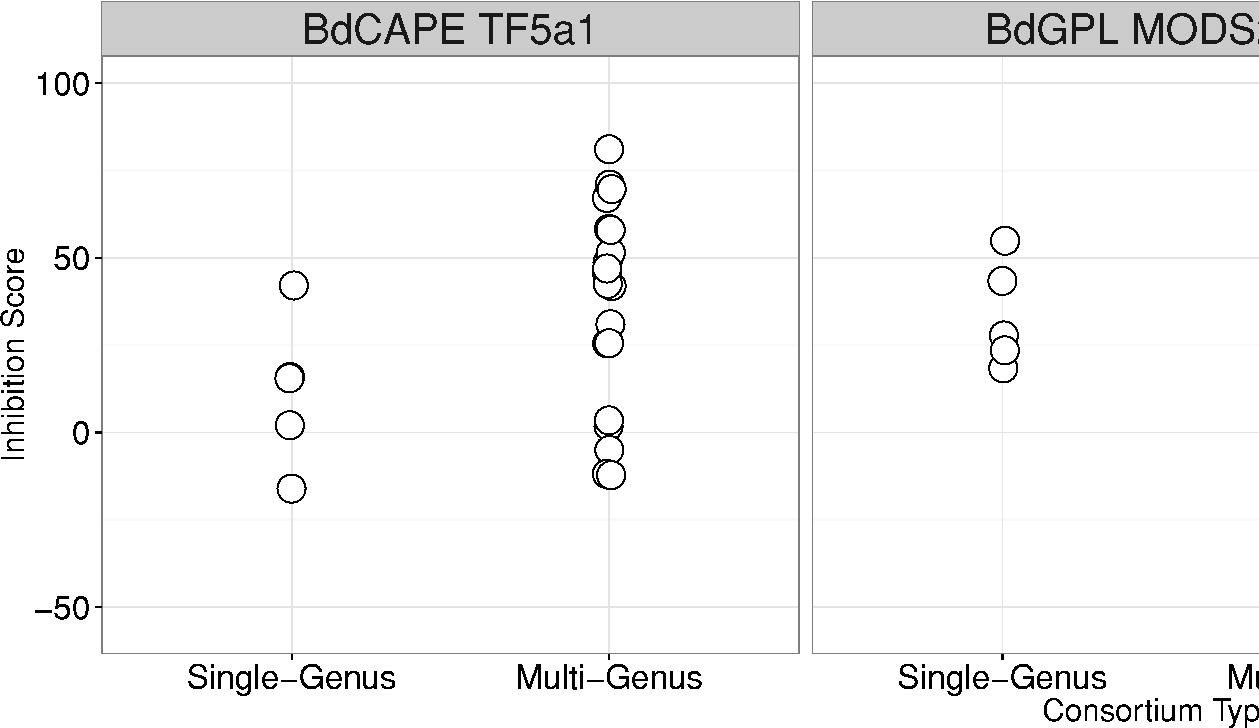
Inhibition scores for Single-Genus and Multi-Genus Consortia across three *B. dendrobatidis* isolates. (*Bd*GPL MODS28.1, *Bd*GPL SFBC019 and *Bd*CAPE TF5a1). Points have been jittered for display purposes.

### in silico Probiotic Consortia Trials

Of the 1000 simulated probiotic trials, naïve application of a MG consortium yielded higher *B. dendrobatidis* inhibition in 69.4% of cases [95% CI 66.5-72.3%] compared to SG consortia (null expectation 50%, p_RAND_<0.001). Moreover, MG consortia had a 38.1% [35.1 – 41.1%] probability of yielding inhibition greater than 50% (strong inhibition), compared to only 13.9% [11.8 – 16.1%] probability for SG consortia. Mean inhibition for all MG consortia was 36.7%, compared to 16.47% for SG consortia. MG consortia outperformed the single isolate in 61% [58-64%] of cases (null expectation 50%, p_RAND_<0.001). However, by averaging over all *B. dendrobatidis* isolates, these results masked substantial variation among *B. dendrobatidis* isolates in the relative efficacy of MG versus SG consortia. We repeated the above simulations separately for each *B. dendrobatidis* isolate, and found that MG consortia were superior to SG consortia and single bacterial isolates for only two *B. dendrobatidis* isolates (*Bd*GPL MODS28 and *Bd*CAPE TF5a1), and performed slightly worse than SG consortia for *Bd*GPL SFBC019 (Fig. 4A). Moreover, although MG consortia have the greatest probability of yielding >50% inhibition for *Bd*GPL MODS28 and *Bd*CAPE TF5a1, this was not the case for *Bd*GPL SFBC019, where SG consortia had a marginally higher probability of delivering strong inhibition (Fig. 4B).

**Figure 4.**
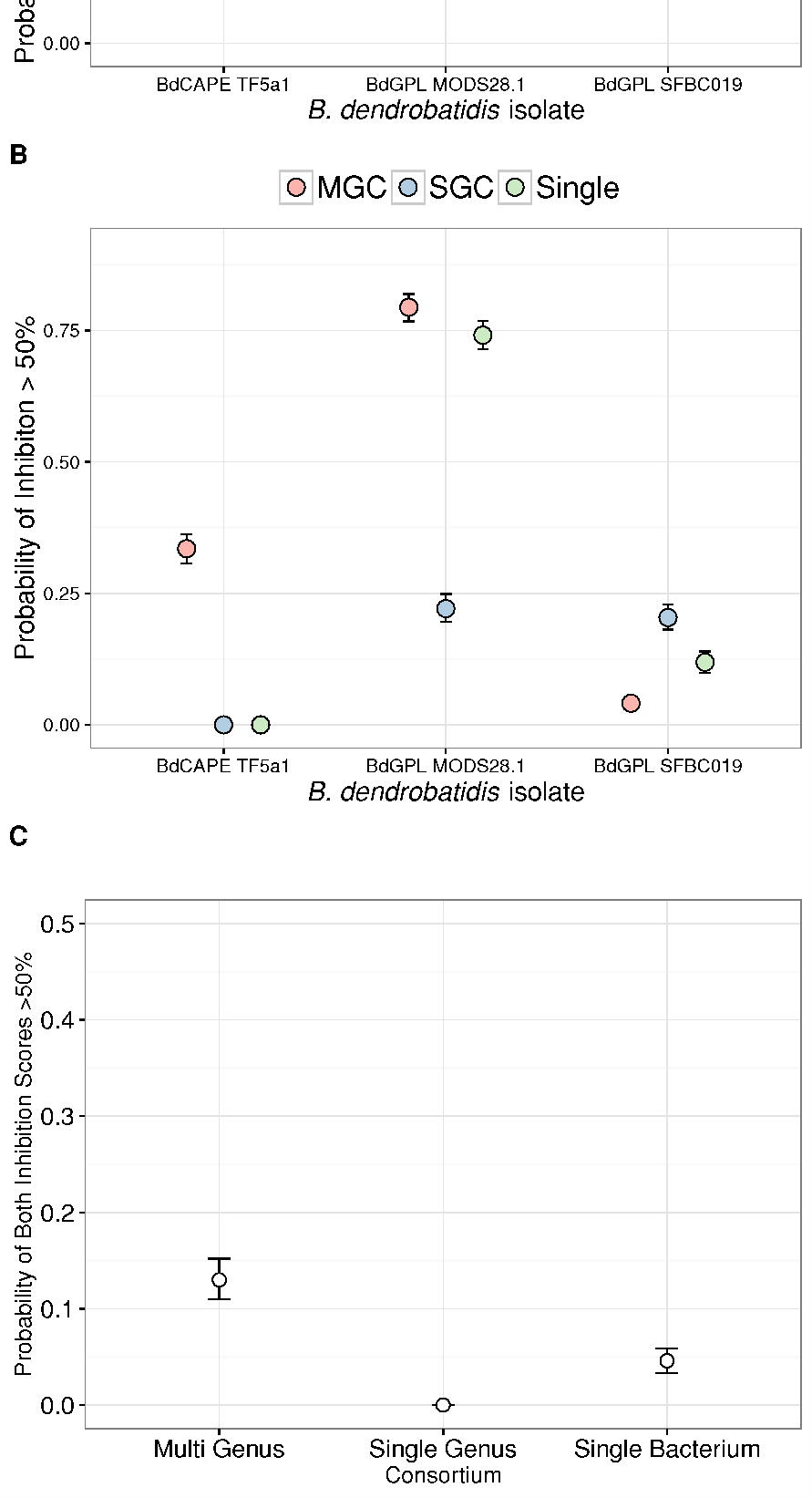
Simulation results examining the relative efficacy of different probiotic strategies. (A) the probability of Multi-Genus Consortia (MGC) yielding higher inhibition compared to Single-Genus Consortia (SGC) or a Single Bacterial Isolate (Single); (B) the probability of MGC, SGC or Single bacteria yielding inhibition > 50% when applied to each of three *B. dendrobatidis* isolates; (C) The probability of an individual consortium type yielding >50% inhibition when applied to two randomly chosen *B. dendrobatidis* isolates in series.

Finally, *we* tested the ability of both MG and SG consortia to inhibit the growth of two different *B. dendrobatidis* isolates in series, as individuals in a single location may be exposed to multiple variants of a pathogen (Goka et al 2009; Schloegel et al 2012; Rodriguez et al 2014; Jenkinson et al 2016), or strong spatial structure of the pathogen and high host dispersal may expose individuals to multiple pathogen variants consecutively. For a given trial, the modelling outcomes were; i) MG consortia inhibited both *B. dendrobatidis* isolates more strongly than SG consortia; ii) SG consortia inhibited both *B. dendrobatidis* isolates more strongly than MG consortia; iii) MG inhibited the first *B. dendrobatidis* isolate more strongly than SG consortia, but not the second; iv) MG inhibited the second *B. dendrobatidis* isolate more strongly than SG consortia, but not the first. Applying the same MG consortium to two *B. dendrobatidis* isolates in series achieved stronger inhibition than SG consortia in 49.4% [46.3 – 52.5%] of cases (i.e. modelling outcome i; null expectation 25% [0.5^2^], p_RAND_<0.001). This compared to only 7.9% [6.4-9.6%] of cases where SG consortia exhibited superior inhibition for both *B. dendrobatidis* isolates (i.e. modelling outcome iv). Mean inhibition across both *Bd* isolates for the MG consortia was 73.4%, compared to 32.5% for SG consortia. MG consortia provided superior inhibition for only one of the *B. dendrobatidis* isolates in the remaining 43% of cases (mean 20.3% and 22.4% of simulations with superior inhibition for the first and second isolate respectively). MG consortia exhibited strong inhibition (>50%) for both isolates in 14.7% [12.5-17%] of cases, compared to zero cases where SG isolates did so. Applying a single bacterial isolate instead of a SG or MG consortium resulted in strong inhibition for both *B. dendrobatidis* isolates in only 4% [2.9-5.3%] of cases (Fig. 4C).

## DISCUSSION

The principal objectives of this study were two-fold: i) to determine the magnitude, if any, of phylogenetic signal in the ability of certain genera of bacteria to inhibit a broad range of *Bd*GPL isolates; and ii) to examine the relative effectiveness of single bacteria and bacterial consortia to inhibit several isolates of *B. dendrobatidis*. We found no evidence of variation among bacterial genera in their ability to exhibit broad-range inhibition across multiple *Bd*GPL isolates. Furthermore, our data suggested consortia provide superior *B. dendrobatidis* inhibition than individual bacteria, but critically this pattern is not uniform across pathogen isolates, and is contingent on consortium taxonomic diversity. Our results have important implications for our understanding of the factors determining *in vivo* resistance to infection in the wild, and provide novel insights into effective strategies for designing probiotic therapies to mitigate lethal cutaneous infections.

### Phylogenetic Signals of Bd GPL Inhibition

We detected no phylogenetic signal in the ability of individual bacterial genera to inhibit multiple *Bd*GPL isolates. These data support previous work suggesting the ability to inhibit *B. dendrobatidis* is distributed widely over bacterial genera (Antwis et al 2015; Becker et al 2015); several isolates demonstrated at least weak inhibition for all 10 *Bd*GPLs but were spread across multiple genera with no clear pattern. That there is clear functional redundancy among genera in this host-protective trait suggests it is not prudent to focus on any one genus in the search for beneficial probiotics (Becker et al 2015), as highly divergent microbial communities can still possess similar functional traits (e.g. Bletz et al 2016). The principal source of variance in inhibition was among bacterial strains, with the number of isolates demonstrating broad-spectrum *facilitation* of *Bd*GPL being roughly equal to the number exhibiting broad-scale *inhibition* of the pathogen. The phenomenon of *Bd*GPL growth facilitation has been described previously for single pathogen isolates (Bell et al 2013; Becker et al 2015), but crucially our results suggest that a bacterial strain’s ability to facilitate the growth of *B. dendrobatidis* may extend across a broad suite of pathogen isolates.

It is unclear why some bacterial isolates facilitate *B. dendrobatidis* growth, but one likely explanation is that certain bacterial metabolites can act as growth substrates for fungi (Garbaye 1994; Hardoim et al 2015), or that different bacterial metabolites alter the abiotic environment (e.g. pH) to confer different growth rates (Romanowski et al 2011). Here we have provided some of the first evidence that facilitation of *B. dendrobatidis* growth is not simply a rare phenomenon arising from specific *Bd*GPL/bacterial combinations, but that this is widespread across bacterial isolates, and different *Bd*GPL isolates differ systematically in their growth rates when exposed to bacterial metabolites. That said, all four CORN isolates showed similar levels of inhibition across all bacterial isolates, whereas the two AUL isolates exhibited markedly different inhibition profiles (Figure 2). We identified one *Bd*GPL isolate that was significantly prone to inhibition, and a further two isolates that demonstrated strong resistance to inhibition across the 58 bacterial isolates we tested. That there is variation in this trait among *Bd*GPL isolates is intriguing; if facilitation occurs because *B. dendrobatidis* uses bacterial metabolites for nutrition, it may suggest some *B. dendrobatidis* variants can use those metabolites more efficiently for growth. Data gathered from additional isolates will allow us to formally test this hypothesis by probing whether a *Bd*GPL’s susceptibility to inhibition or facilitation correlates with virulence. Previous work has shown no among-isolate variation in susceptibility of *B. dendrobatidis* to an echinocandin antifungal drug (Fisher et al 2009), yet our data suggest this pattern is not the same for bacterial metabolites. Recombination among lineages of *Bd*GPL is common (Farrer et al 2011), providing a mechanism whereby metabolic genes favouring enhanced growth may be spread following contact among lineages. Our data have two important implications given the proclivity of *B. dendrobatidis* for recombination. First, among-isolate variation in susceptibility to inhibition suggests that the relative efficacy of probiotic or curative therapies in the wild will be modified by local *B. dendrobatidis* genotype. Second, though we tend to treat bacterial inhibition scores as fixed traits, this ignores the ability of genetic recombination among *B. dendrobatidis* lineages to modify the relationship between bacterial metabolites and pathogen growth rates. Even the application of probiotics themselves may represent a strong selective pressure favouring genetic variants of *B. dendrobatidis* that lack susceptibility to those probiotics. Although several trials have demonstrated the potential for probiotic prophylaxis against *B. dendrobatidis* (e.g. Harris 2009; Muletz et al 2012; Loudon et al. 2014; Kueneman et al 2016), we still lack the requisite data to measure selection caused by those trials on the pathogen. *In vitro* experimental evolution assays between pathogen and bacteria may prove the most powerful means for detecting such patterns.

### Consortium-Based Approaches to Combatting Fungal Pathogens

Our results revealed a positive link between the taxonomic richness of a probiotic consortium and its ability to inhibit *B. dendrobatidis* growth, but crucially this relationship was highly dependent on *B. dendrobatidis* isolate. Multi-genus consortia outperformed both single-genus consortia and single bacterial isolates in *B. dendrobatidis* inhibition, and were far more likely to produce strong inhibition of 50% or greater, but only for two of the three pathogen variants.

The general relationship between inhibition and consortium diversity was in the expected direction; low community relatedness (i.e. high community dissimilarity) and high species richness both increase the resistance of a bacterial community to pathogenic ‘invaders’ (e.g. Jousset et al 2011; Eisenhauer et al 2012, 2013). Furthermore, previous work has linked higher species diversity of probiotic consortia to increased *B. dendrobatidis* inhibition using a single pathogen isolate (Loudon et al 2014; Piova-Scott et al 2017). Superior inhibition from consortia, rather than single isolates, may arise as a by-product of the interference competition over resources created by co-culture (Scheuring & Yu 2012). Thus, even bacteria that are weak inhibitors when grown individually can increase the overall inhibitory power of a consortium by creating a competitive environment that favours greater production of anti-fungal compounds. Functional dissimilarity has been proposed as more important than taxonomic diversity in predicting a community’s resilience to invasion (Eisenhauer et al 2013), but may explain why single-genus consortia did not perform as well as multi-genus consortia. In selecting for genetic diversity, we may have been simultaneously selecting for functional diversity not present when co-culturing three members of the same genus.

That *B. dendrobatidis* isolate can alter the strength of the relationship between consortium diversity and inhibition is a highly novel finding. Our simulated probiotic trials revealed that for two *B. dendrobatidis* isolates, combining bacteria into multi-genus consortia yielded significantly better inhibition than applying one of the member bacteria in isolation. These results provide further support for a synergistic effect of co-culture on inhibition. If multi-genus consortia were no better at inhibition than the mean of their composite members, Monte Carlo integration over all single isolate scores would not have recovered a significant difference between the two groups. Yet, for *Bd*GPL MODS28 and *Bd*CAPE TF5a1, multi-genus consortia yielded by far the highest probability of observing strong inhibition of 50% or more. That this pattern was not conserved for *Bd*GPL SFBC019 is perhaps the most intriguing finding. As for *Bd*GPL variants JEL423 and AUL2 in the phylogenetic trials, SFBC019 was largely resistant to inhibition, with individual bacterial inhibition scores that were often negative. One possible explanation for the lack of efficacy of consortia against SFBC019 is that the when a variant of *B. dendrobatidis* is resistant to inhibition and/or there is little variation in inhibition, co-culture fails to produce any synergistic inhibitory effects. That is, if a pathogen is highly resistant to most bacterial metabolites in the first instance, increases in the relative concentrations of those metabolites through co-culture-mediated competition are unlikely to elicit any significant increases in inhibitory capability. The most important consequence of this pattern is that for some pathogenic variants, multi-genus consortia are unlikely to be able to yield high inhibition in cases where individual isolates have failed to do so. Despite the observed variance in success of consortia across *B. dendrobatidis* isolates, our simulation trials revealed that multi-genus consortia offer the best broad-spectrum protection across multiple *B. dendrobatidis* isolates encountered in series. This finding is important; human-mediated spread of *B. dendrobatidis* through the amphibian trade (Fisher & Garner 2007) means we cannot assumise that local populations will be exposed to only one pathogenic variant. Successful mitigation of the pathogen in the wild demands that we employ strategies with the highest broad-spectrum success over multiple pathogen genotypes. Combining bacteria that show high levels of inhibition across multiple *B. dendrobatidis* isolates may further increase the effectiveness of consortia (intermediate inhibitors were selected for this study to allow greater insight into community dynamics).

### Conclusion

This study adds to a growing body of evidence suggesting that diverse, multi-species consortia may represent powerful disease mitigation tools, offering superior probiotic protection against disease compared to single bacterial isolates. Our work has highlighted that different isolates of a pathogen can modify the strength of inhibition caused by the probiotic, meaning we cannot expect probiotic effectiveness to be uniform across the genetic landscape of the pathogen. Despite the relative merits of multi-genus consortia for mitigating single and multiple *B. dendrobatidis* variants, it remains to be determined how readily these consortia will be able to colonise the host skin *in vivo*. This is crucial for to being able to quantify how applicable inhibition measures derived *in vitro* are to real-world scenarios. Nevertheless, our data highlight the merits of a community-level approach to probiotic mitigation of wildlife disease, which may offer more broad-spectrum host protection in the face of large-scale heterogeneity in pathogen genotype.

## ACKNOWLEDGEMENTS

This study was partly funded by a North-West University Postdoctoral Research Fellowship awarded to REA. XAH was funded by an Institute of Zoology Research Fellowship. The authors would like to thank Prof Richard Preziosi and Dr Trenton Garner for additional provision of consumables, and Prof Ché Weldon, Dr Trenton Garner and Prof Matthew Fisher for access to *Batrachochytrium dendrobatidis* isolates used in this study.

## DATA ACCESSIBILITY

Genbank accession numbers of all bacteria are provided in Supplementary Materials. Data files and R Markdown will be made available on Dryad.

## AUTHOR CONTRIBUTIONS

RA and XH conceived the study, RA collected the data, XH analysed the data, RA and XH wrote the paper. Both authors contributed equally to this paper.

## TABLE LEGENDS

**Table 1**

*Batrachochytrium dendrobatidis* isolates used in the study.

**Table 2**

Composition of multi-genus consortia used in the study. Single-genus consortia comprised all three bacterial isolates (A, B and C) for a given genus (*Acinetobacter, Chryseobacterium, Serratia, Stentrophomonas*).

**Table 3**

Mean Proportion of 10 *Bd*GPL isolates for which at least weak inhibitory capability was observed, averaged over all bacterial isolates in a genus. 95% CI: 95% confidence intervals from an overdispersion-corrected Binomial GLM.

## Notes

**Conflict of Interest:** The authors declare no conflict of interest.

